# The copper resistome of group B *Streptococcus* reveals insight into the genetic basis of cellular survival during metal ion stress

**DOI:** 10.1101/2021.09.15.460573

**Authors:** Kelvin G. K. Goh, Matthew J. Sullivan, Glen C. Ulett

## Abstract

In bacteria, copper (Cu) can support metabolic processes as an enzymatic cofactor but can also cause cell damage if present in excess, leading to intoxication. In group B *Streptococcus* (GBS) a system for control of Cu efflux based on the canonical *cop* operon supports survival during Cu stress. In some other bacteria, genetic systems additional to the *cop* operon are engaged during Cu stress and also contribute to Cu management. Here, we examined genetic systems beyond the *cop* operon in GBS for regions that contribute to survival of GBS in Cu stress using a forward genetic screen and probe of the entire bacterial genome. A high-density mutant library, generated using pGh9-IS*S1*, was used to expose GBS to Cu stress and compared to non-exposed controls *en masse*. Nine genes were identified as essential for GBS survival in Cu stress, whereas five genes constrained GBS growth in Cu stress. The genes encode varied factors including enzymes for metabolism, cell wall synthesis, transporters and global transcriptional regulators. Targeted mutation of the genes validated their roles in GBS resistance to Cu stress. Notably, several genes, including *stp1, yceG, plyB* and *rfaB* were also essential for resistance to Zn stress. Excepting *copA*, the genes identified are new to the area of bacterial metal ion intoxication. We conclude that a discrete and limited suite of genes beyond the *cop* operon in GBS contribute to a repertoire of mechanisms used to survive Cu stress *in vitro* and achieve cellular homeostasis.

**Significance Statement:** Genetic systems for copper (Cu) homeostasis in bacteria, including Streptococci, are vital to survive metal ion stress. Genetic systems that underpin survival of GBS during Cu stress, beyond for the archetypal *cop* operon for Cu management, are undefined. We show that *Streptococcus* resists Cu intoxication by utilizing a discrete and limited suite of genes beyond the *cop* operon, including several genes that are new to the area of bacterial cell metal ion homeostasis. The Cu resistome of GBS defined here enhances our understanding of metal ion homeostasis in GBS.

## Introduction

In prokaryotic and eukaryotic cells, copper (Cu) is an important cofactor for metalloenzymes (1). When present in excess however, Cu can cause cytotoxicity. In bacteria, Cu intoxication can reflect enzyme inactivation, deregulation of metabolism, and/or redox stress, including a higher potential to generate reactive oxygen species (2). In the context of an infected host, phagocytes such as macrophages and neutrophils can mobilise intracellular pools of Cu to pro-actively expose internalized bacteria to excess metal to achieve conditions that are antimicrobial (3, 4). Such subcellular areas within infected phagocytes in which concentrated Cu exerts antimicrobial effects have been described for several bacterial pathogens (5, 6). The antimicrobial properties of Cu are thus of interest to the field of infection and immunity since these offer potential avenues for antimicrobial benefit, which might be harnessed to better control bacterial infection (4, 7, 8).

The canonical system for Cu efflux in bacteria utilizes the *cop* operon, encompassing *copA* that encodes an ATPase efflux pump that extrudes cellular Cu ions, alongside a Cu-specific transcriptional regulator *copY*, that represses the operon (9-11). Adaptation to metal excess and limitation in bacteria is nonetheless complex. Several systems additional to the *cop* operon based on efflux proteins, including P-type ATPases are also described, and these contribute to bacterial resistance to metal stress for several pathogens, as reviewed elsewhere (5). *Streptococcus agalactiae*, also known as group B *Streptococcus* (GBS) is an opportunistic bacterial pathogen of humans and animals for which a discrete genetic system for cellular management of Cu homeostasis based on the *cop* operon was recently described (12). The GBS *cop* operon regulates Cu import and export by responding to excess Cu and de-repressing *copA* via CopY to drive Cu export from the cell (12). Transcriptional activation of the *cop* operon in GBS in response to Cu stress was also shown to contribute to virulence of the bacteria during acute infection (12).

Here, we sought to identify genetic systems additional to the *cop* operon that aid Cu management in GBS. We used a genome-wide approach based on Transposon Directed Insertion Site Sequencing (TraDIS) to probe the GBS genome for regions that support cell survival of Cu stress.

## Results

### Determination of growth conditions and Cu concentration required for TraDIS

To probe the entire GBS genome for regions that support the survival of this organism during Cu stress, we first evaluated the conditions for *in vitro* exposure of GBS to Cu stress, which would be suitable for a subsequent forward genetic screen. To do this, we tested a Cu concentration of 1.5 mM in Todd-Hewitt Broth (THB) medium for inhibition of GBS growth because this was sufficient to inhibit a GBS mutant deficient in *copA* (encoding a Cu efflux P-type ATPase) in a prior study (12). Growth assays verified that 1.5 mM Cu in THB was insufficient to inhibit the growth of wild-type GBS 874391 but sufficient to completely inhibit the growth of a GBS Δ*copA* mutant (Fig.1). The level of 1.5 mM Cu in THB was therefore accepted as suitable to probe for additional genes that contribute to resistance to Cu stress, as this would inhibit growth of mutants sensitive to Cu.

**Figure 1.**
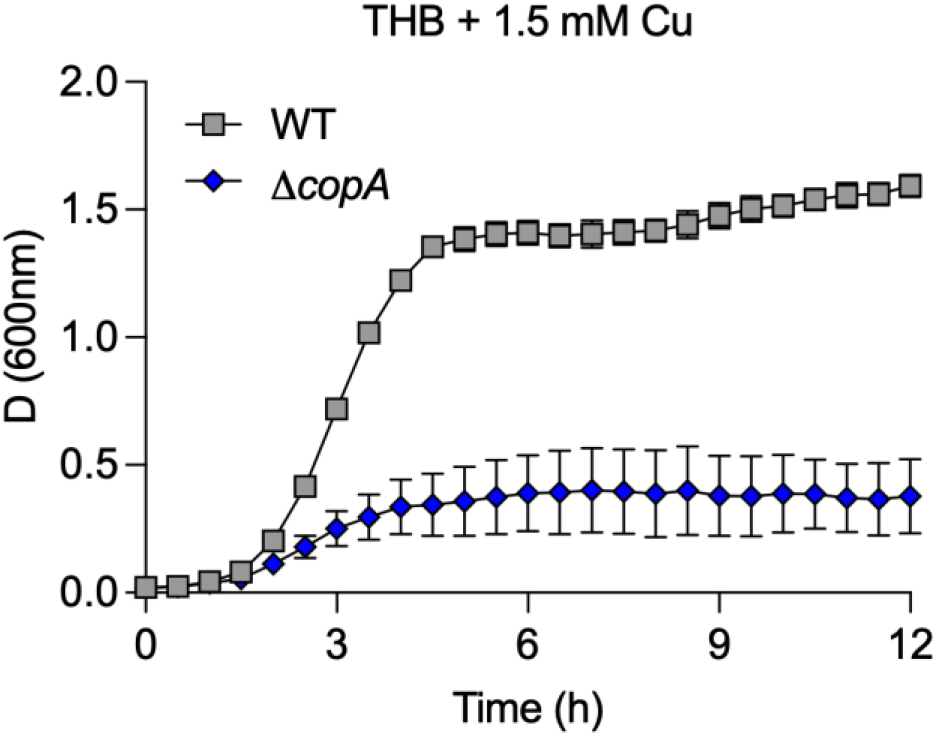
Growth of WT GBS 874391 and a *copA*-deficient mutant in Cu stress. The bacteria were grown in THB supplemented with 1.5 mM Cu for 12 h. Points show means of attenuance (600nm) and bars show s.e.m. (*n*=3).

### Identification of genes associated with Cu resistance in *GBS* 874391

To facilitate an extensive genome wide screen of genes required for Cu resistance by 874391, we first generated a library of approximately 480,000 mutants by using a pGh9-IS*S1* plasmid carrying an erythromycin resistance gene. Next, we subjected the mutant library to the growth conditions we established earlier to identify genes associated with Cu resistance. In this assay, roughly 1.9 × 10^8^ cells from the library were inoculated in triplicate into 100ml of THB either supplemented with 1.5mM Cu (test pool) or without supplementation of Cu (control pool), and incubated for 12 h at 37°C. This permitted the growth of mutants unaffected by Cu, but inhibited growth of mutants sensitive to Cu. Genomic DNA was extracted from each replicate and sequenced with a multiplex TraDIS approach (Fig. 2A). The control pool yielded a total of 5,163,397 IS*S1-* specific reads that mapped to the 874391 genome. Further analysis of the control data revealed 618,263 unique insertion sites (approximately one insertion site every four bp) distributed across the entire chromosome (Supplementary Fig. S1), highlighting the high level of insertion saturation and coverage of our library.

**Figure 2.**
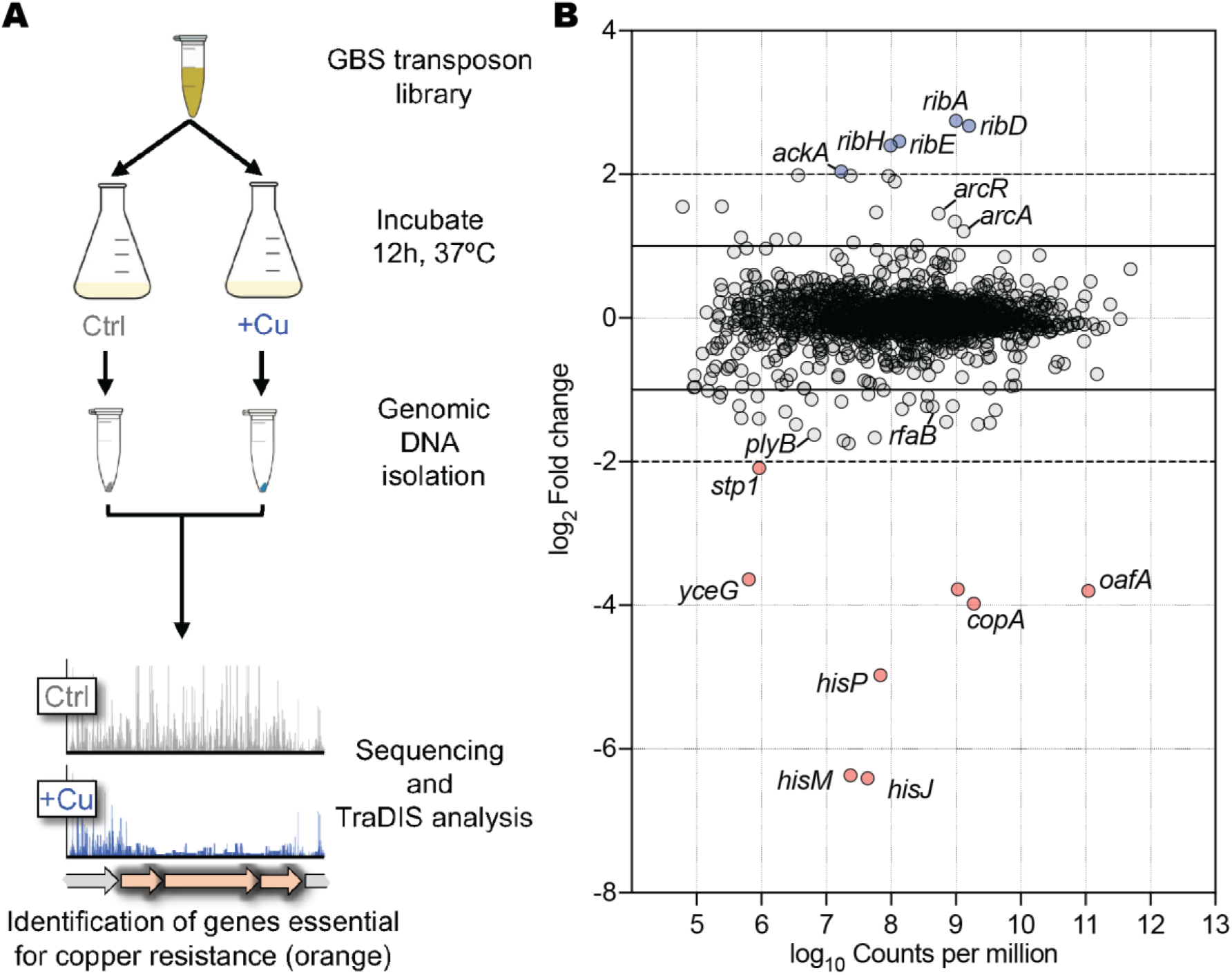
Defining the Cu resistome of GBS using TraDIS. (A) Experimental design to identify genes associated with Cu sensitivity. Approximately 1.9 × 10^8^ cells were inoculated into THB or THB + 1.5 mM Cu and grown for 12 h at 37°C with agitation at 200rpm. Bacterial genomic DNA was then purified and subjected to TraDIS analysis. (B) TraDIS analysis of the GBS Cu resistome identified 5 genes that were significantly over-represented (blue), and 8 genes that were significantly under-represented (red), using highly stringent cut-offs (2 < log_2_FC< -2; FDR <0.001 and P value <0.05). A further 28 and 15 were significantly under-or over-represented between 2-4-fold, respectively. Horizontal solid lines highlight FC cutoffs of 2 < log_2_FC< -2 and grey dashed lines indicate cutoffs of 1 < log_2_FC< -1.

As mutants with insertions in genes required for Cu resistance would be lost and underrepresented, we screened for a loss of insertions in the test pool as compared to the control pool. Using a stringent selection criteria of a log_2_ fold change (log_2_FC) in read counts of ≤ -2, false discovery rate (FDR) of <0.001 and P value of <0.05, we identified a hit plot of genes that contributed to GBS growth in Cu stress (Fig. 2B). Here, we identified nine genes as highly significantly under-represented in the dataset (Table 1). Consistent with previous reports of a requirement for *copA* in resisting Cu stress, *copA* was significantly under-represented (∼16-fold down) during Cu stress, representing validation for the TraDIS approach. Interestingly, we also identified five genes that possessed an enrichment of read counts (log_2_FC of ≥ 2) in the test pool compared to the control pool, suggesting that insertions in these genes were beneficial for growth under Cu stress (Table 1). Representative insertion site mapping is shown for a selection of loci (Fig. 3A-F). We also noted a further 28 and 15 genes that were under-represented and over-represented in the dataset (log_2_FC between -1 to -2 or 1 to 2; Supplementary Table S1), respectively. Interestingly, this list included two genes (*rfaB* [log_2_FC = -1.24] and *plyB* [log_2_FC = -1.63]) which we previously found to be essential for zinc resistance (13). Hence, we decided to also include these two genes in our further analyses.

**Table 1.** Genes identified by TraDIS that play a role in Cu sensitivity in 874391.

| Locus tag<sup>#</sup> | Gene | Annotation | log<sub>2</sub>FC | log<sub>10</sub>CPM |
| --- | --- | --- | --- | --- |
| 01047 | *hisM* * | amino acid ABC transporter permease | -6.41 | 7.63 |
| 01048 | *hisJ* * | amino acid ABC transporter ATP-binding protein | -6.37 | 7.37 |
| 01049 | *hisP* * | amino acid ABC transporter substrate-binding protein | -4.94 | 7.98 |
| 00507 | *copA* * | Cu-translocating P-type ATPase | -3.98 | 9.27 |
| 00084 | *oafA* * | acyltransferase | -3.80 | 11.04 |
| 00083 | - | membrane protein | -3.78 | 9.02 |
| 01646 | *yceG* * | MltG-like endolytic transglycosylase | -3.64 | 5.80 |
| 00435 | *stp1* * | Stp1/IreP family PP2C-type Ser/Thr phosphatase | -2.10 | 5.96 |
| 00288 | *ackA* | acetate kinase | 2.04 | 7.22 |
| 00879 | *ribH* | 6 and 7-dimethyl-8-ribityllumazine synthase | 2.40 | 7.98 |
| 00877 | *ribE* | riboflavin synthase | 2.46 | 8.12 |
| 00876 | *ribD* * | bifunctional diamino-hydroxyphospho-ribosyl-amino-pyrimidine deaminase/5-amino-6-(5-phosphoribosylamino) uracil reductase RibD | 2.67 | 9.19 |
| 00878 | *ribA* | bifunctional 3 and 4-dihydroxy-2-butanone-4-phosphate synthase/GTP cyclohydrolase II | 2.74 | 9.00 |

<sup>#</sup>denotes GBS str 874391 locus tag, preceded by CHF17\_ ; \* denotes genes that were mutated

**Figure 3.**
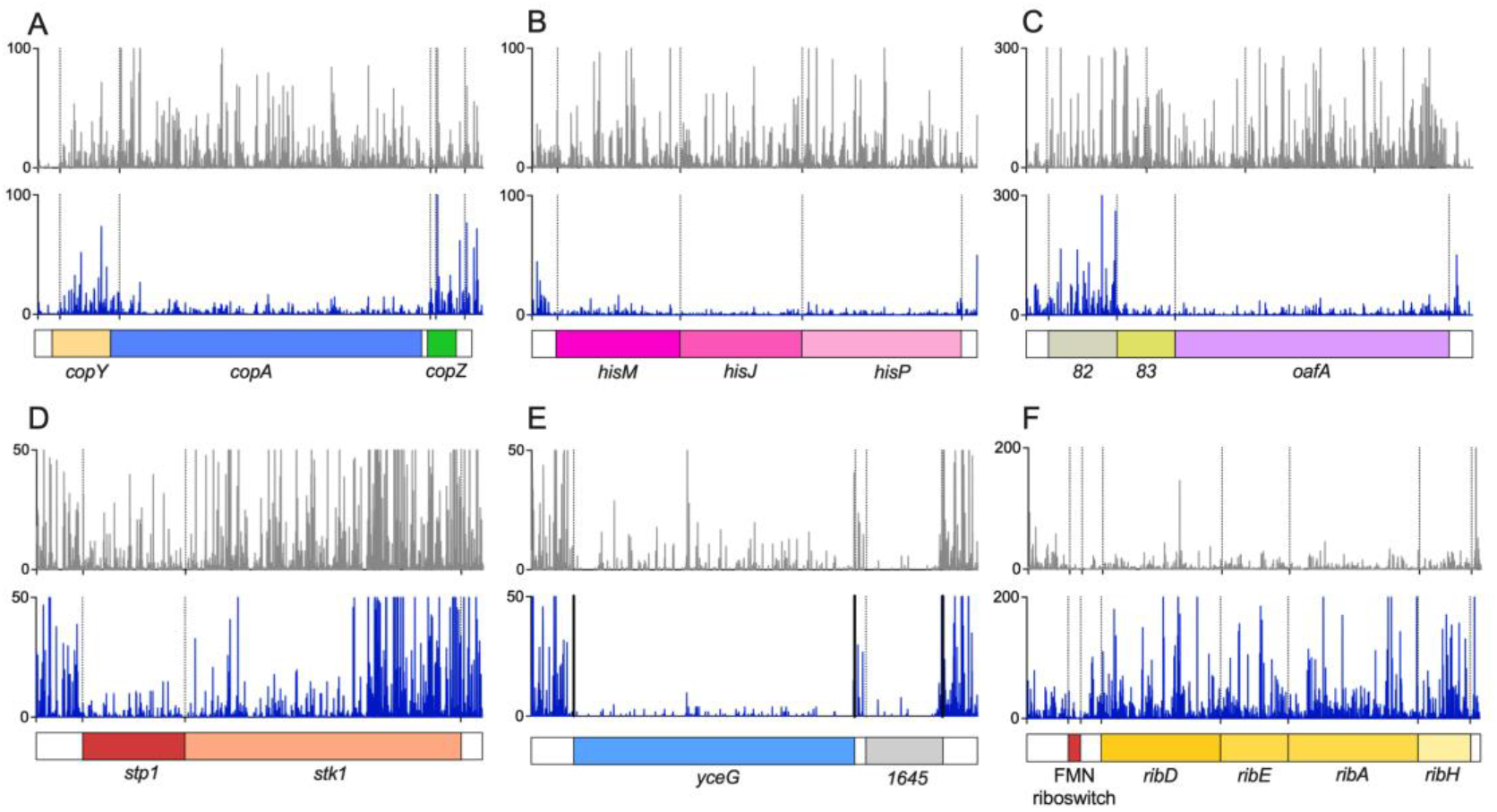
Insertion plots of genes associated with Cu resistance as identified by TraDIS. Individual insertions mapped to the *copYAZ* (A), *hisMJP* (B), *oafA* (C), *stp1* (D), *yceG* (E) and *ribDEAH* (F) loci are shown, with vertical dotted lines denoting the boundaries of each genetic element. Data from the non-exposed control are shown in grey, with reads from the Cu-stress condition shown in blue.

### Characterisation and validation of Cu sensitive mutants

To validate hits from the TraDIS screen, we generated targeted isogenic mutants of several candidate genes and phenotypically analysed these mutants for survival and growth in Cu stress. The genes included *hisMJP* (CHF17_01047, 01048 and 01049), *oafA* (CHF17_00084), *yceG* (CHF17_01646), *stp1* (CHF17_00435), *ribD* (CHF17_00876), *rfaB* (CHF17_00838), and *plyB* (CHF17_00885). Firstly, colony forming unit (CFU) assays based on the conditions used for the TraDIS screen were performed to test survival of the mutants in 1.5 mM Cu after a 12 h incubation period. In this assay, besides WT and the Δ*hisMJP* mutant, all other isogenic mutants exhibited a significant decrease (∼30 to 72%) in CFU counts when grown in the presence of Cu, indicating that these mutants were sensitive to Cu (Fig 4A). Notably, we also observed a significantly lower overall bacterial counts in the Δ*plyB* and Δ*stp1* mutants as compared to WT when grown in THB, indicating that these genes contribute to the growth of GBS in rich media.

**Figure 4.**
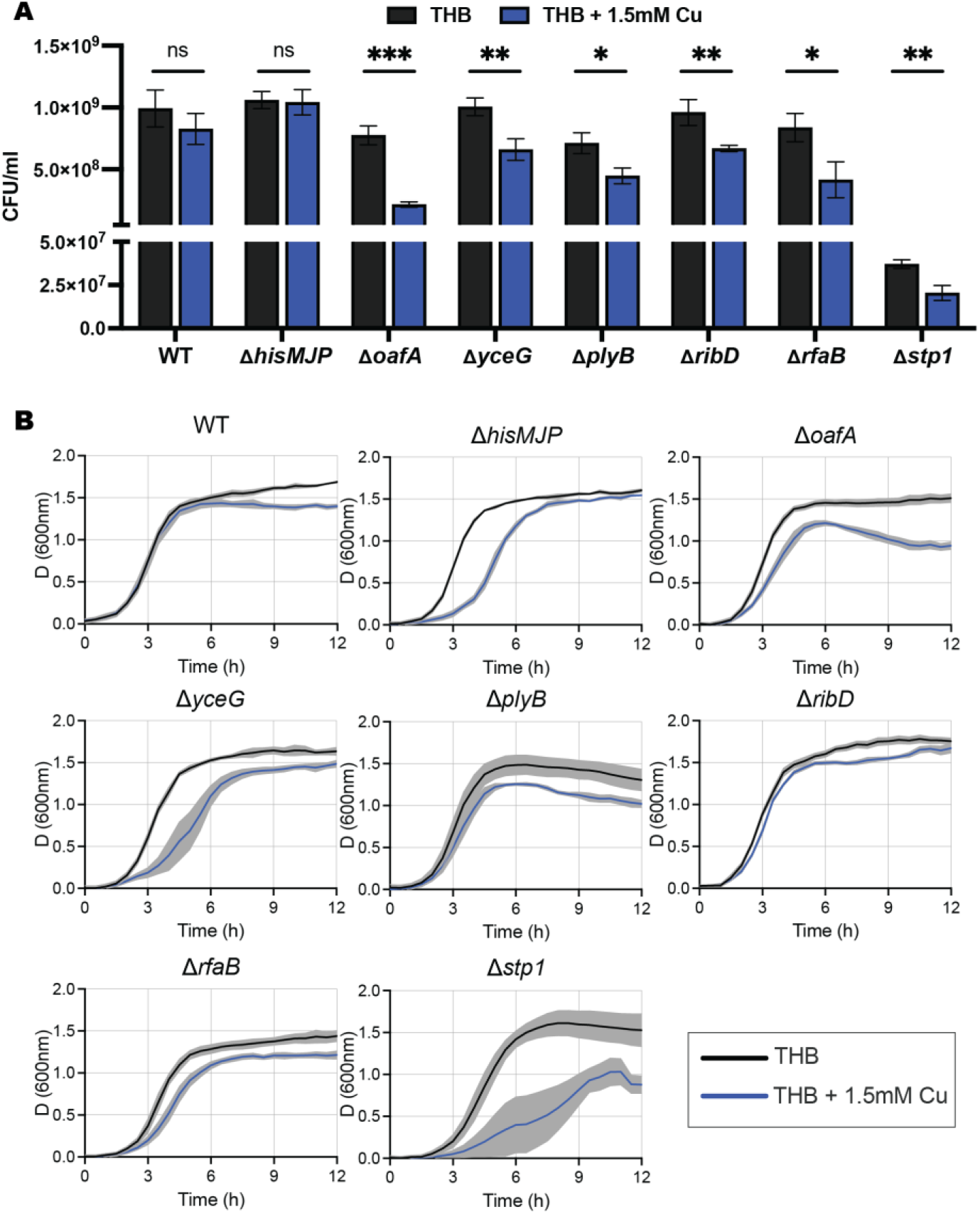
Novel GBS genes that contribute to resistance to Cu stress in THB media. (A) Colony forming unit (CFU) assays of WT GBS and isogenic mutants grown in THB or THB +1.5mM Cu for 12 h. P values calculated with a pairwise Student’s *T*-test comparing THB and THB + 1.5 mM Cu (***, P < 0.001; **, P < 0.01; *, P < 0.05; ns = not significant). Data are means of 3 independent repeats with error bars indicating SD. (B) Growth curves of GBS in THB with (1.5mM Cu) or without Cu stress. Data are means of ≥ 3 independent repeats with shaded area indicating s.e.m.

We further explored the Cu sensitivity of the mutants in THB by measuring growth rate over 12 h. In this assay, the Δ*hisMJP*, Δ*oafA*, Δ*yceG*, Δ*plyB*, Δ*rfaB* and Δ*stp1* mutants displayed attenuation of growth rate either by exhibiting a lower overall absorbance or an increased lag phase when subjected to Cu stress (Fig 4B). The Δ*ribD* mutant was able to grow better under Cu stress than compared to wild type in this assay, supporting our TraDIS analysis which identified that insertions in the gene enhanced tolerance to Cu stress.

We previously reported that culture media can affect GBS sensitivity to Cu (12). Consequently, we measured the growth of WT GBS to each isogenic mutant in nutrient limited conditions by using a minimal Chemically-Defined Medium (CDM) supplemented with 1.0 mM Cu. Notably, we detected attenuation in the growth of all mutants except Δ*ribD* in CDM supplemented with Cu compared to WT (Fig. 5). Some strains also exhibited growth defects in our assays in the absence of Cu (Δ*hisMJP*, Δ*plyB*, Δ*yceG* and Δ*stp1*), indicating that some of the target genes contribute to the growth of GBS in CDM. Taken together, our data show that the majority of candidate genes identified in our TraDIS screen contribute to Cu sensitivity in GBS strain 874391 in THB, and that this effect is amplified in minimal CDM.

**Figure 5.**
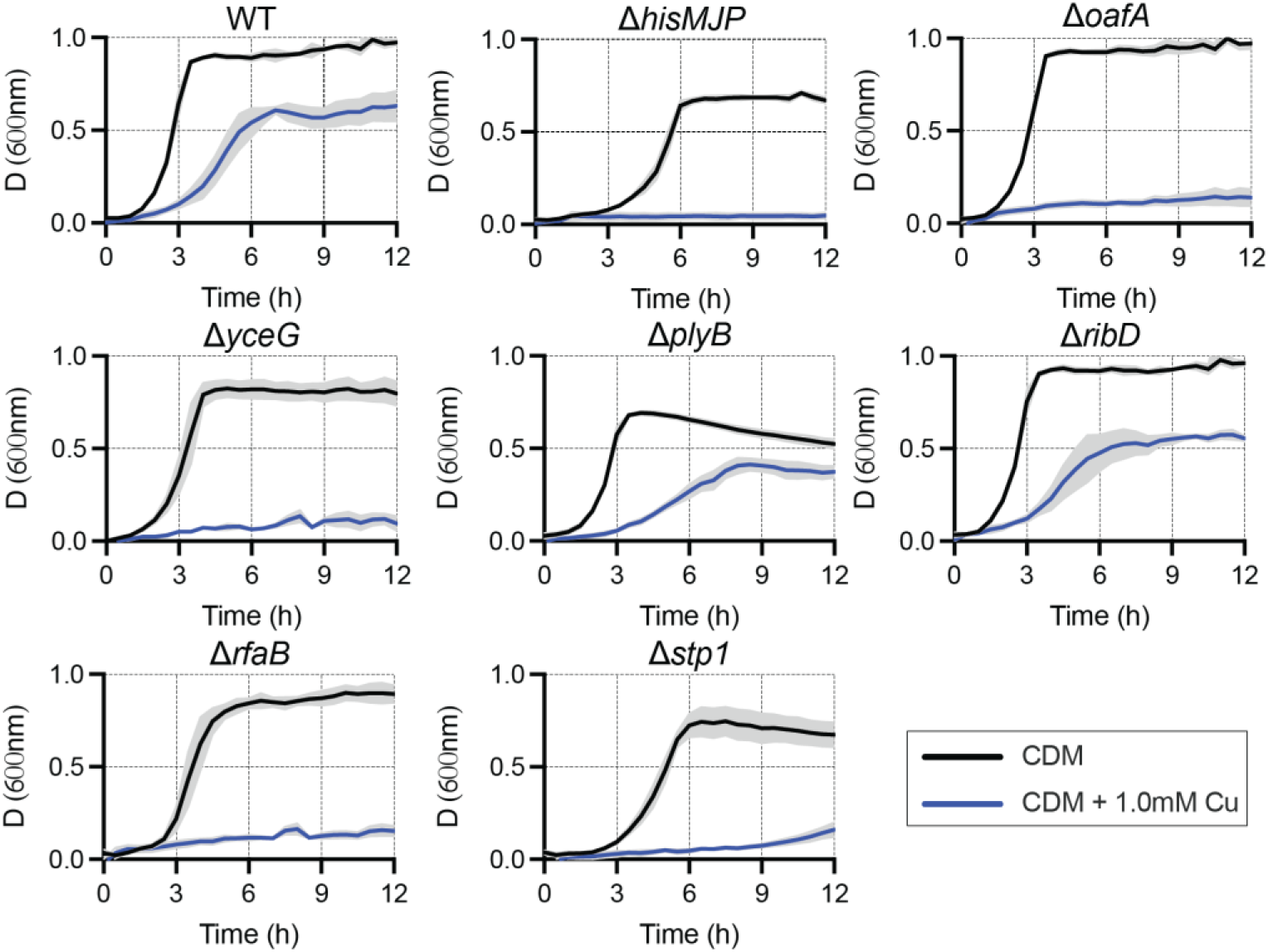
Novel genes involved in resisting Cu stress in minimal CDM. Growth curves of WT GBS and isogenic mutants in CDM supplemented with (1.0 mM) or without Cu. Data are means of ≥ 3 independent repeats with shaded area indicating s.e.m.

### Accumulation of intracellular Cu during Cu stress

Inductively coupled optical emission spectroscopy (ICP-OES) was used to investigate if Cu ion transport is affected in each respective isogenic mutant. Standard THB medium contains 0.2±0.08 μM Cu, reflecting trace amounts in the medium (12). In the absence of supplemental Cu, WT GBS limited intracellular Cu content such that only 0.53±0.04 μg Cu g dry weight^-1^ were detected in cultures grown in THB. Exposure of WT GBS to 1.5 mM Cu resulted in a significant increase in intracellular Cu to 11.69±3.52 μg Cu g dry weight^-1^ (Fig. 6A). This was reflected in the different mutants, where exposure to Cu also resulted in a significant increase in intracellular Cu (Fig. 6). Strikingly, the Δ*stp1* mutant exhibited almost two times more intracellular Cu as compared to WT (20.89±5.85 ug Cu G dry weight^-1^), whereas the Δ*hisMJP*, Δ*ribD* and Δ*rfaB* mutants contained approximately half the intracellular Cu as compared to WT (4.34±1.57, 5.89±1.68 and 6.61±0.77 ug Cu G dry weight^-1^ respectively) (Fig. 6B). On the other hand, the Δ*yceG*, Δ*plyB* and Δ*oafA* mutants all contained comparable intracellular Cu to WT. Taken together, these data indicate that not all the genes identified affect Cu transport, and suggests that sensitivity to Cu in some mutants may not be due to the accumulation of intracellular Cu but through other mechanisms.

**Figure 6.**
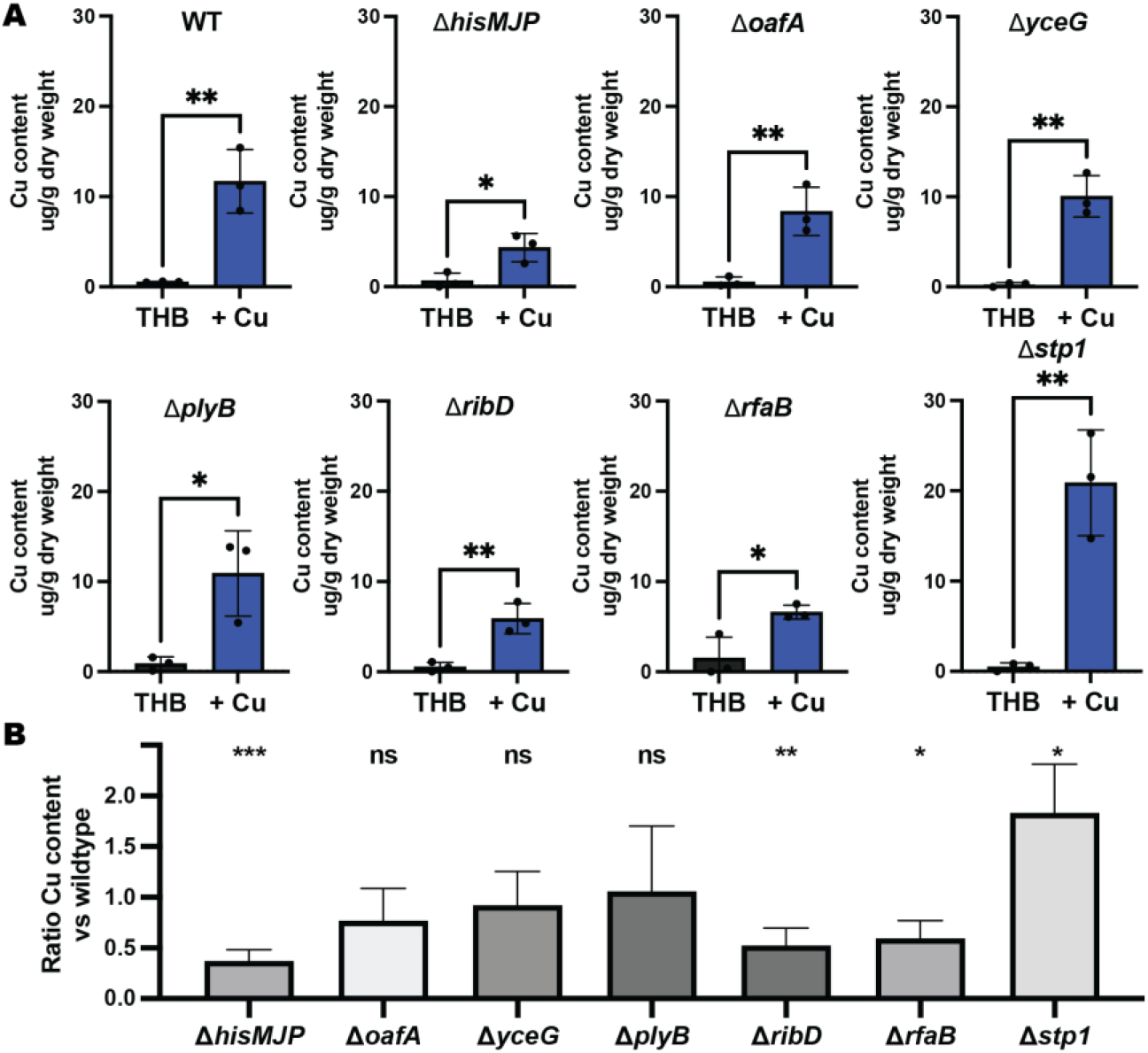
Intracellular Cu content in WT GBS 874391 and isogenic mutants in Cu stress. (A) Total intracellular Cu content was compared in WT and isogenic mutants grown in THB with and without supplemental Cu. P values calculated with a pairwise Student’s *T*-test is presented on the graphs. (B) Ratio of intracellular Cu content in isogenic mutants compared to WT in Cu stress. P values calculated with a pairwise Student’s *T*-test comparing WT to isogenic mutant are presented on top of each bar graph. ***, P < 0.001; **, P < 0.01; *, P < 0.05; ns = not significant. Data presented are means of 3 independent repeats with error bars indicating standard deviation.

## Discussion

GBS is an opportunistic pathogen that causes a diverse range of disease aetiologies in infants and adults, including skin and soft-tissue infections, arthritis, pneumonia, meningitis, urinary tract infection, and endocarditis (14). GBS expresses several virulence factors that enable the bacteria to survive in harsh conditions, such as acid stress, oxidative stress, and during infection of a host, as reviewed elsewhere (15). Metal ion stress due to excess Cu was recently demonstrated to be antimicrobial towards GBS (12). Cu is an essential micronutrient for bacteria (16), but as excess Cu can be toxic to cells, bacteria need to able to regulate the amount of intracellular Cu during Cu stress. This can generally be done using three mechanisms – (i) export of intracellular Cu into the extracellular milieu, (ii) sequestration of Cu by Cu binding proteins, and (iii) oxidation of Cu(I) to the less toxic form of Cu(II) (17). In this study, using a forward genetic screen based on TraDIS, we identify new GBS factors that contribute significantly to the survival and growth of this pathogen in Cu stress conditions. The key findings are that (i) GBS utilizes several genes additional to the *cop* operon to manage Cu homeostasis during Cu stress, (ii) the GBS Cu stress resistome comprises principally nine genes that are required for GBS to resist Cu stress, including *hisMJP, oafA, yceG, plyB, ribD, rfaB* and *stp1*. These genes have not previously been linked to mechanisms of bacterial resistance to Cu stress.

As a screening approach to identify novel functions of bacterial genes, TraDIS has been used to explore GBS survival in blood, which revealed protective effects of calprotectin (18-20). TraDIS analysis in the current study identified novel functions of several genes in GBS that play a role in Cu resistance. We validated our approach by identifying *copA* in our TraDIS screen, which encodes an exporter known to be essential for GBS Cu resistance (12). We generated defined mutants for nine other genes (with Δ*hisMJP* generated as a single mutant). Deletion of these genes resulted in the recovery of lower bacterial counts or attenuated growth rates when compared to the WT under Cu stress, confirming the role of these genes in promoting resistance to Cu. However, the exact mechanism by which these genes facilitate resistance to Cu toxicity is unclear. Four proteins (OafA, RfaB, PlyB and YceG) possess domains commonly found in proteins involved in cell wall biogenesis. Furthermore, the PlyB protein shares structural similarity to a glycosyl hydrolase of *S. pneumoniae*, which was shown to also facilitate the colonisation and invasion of host epithelial cells (21, 22).

Stp1 is a serine/threonine phosphatase and is important for regulation of its cognate kinase Stk1 and GBS virulence (23, 24). We found that not only is the *stp1* mutant more sensitive to Cu stress, but it was also the only mutant that had approximately two times more intracellular Cu as WT while under Cu stress. In GBS, mutation of *stp1* leads to increased phosphorylation of a number of proteins, which in turn affects gene expression. Genes that are downregulated in a *stp1* mutant include several transcriptional regulators and ABC transporters implicated in the uptake of amino acids and metal transport (23). This could explain the Cu sensitivity phenotype we observed in the *stp1* mutant, although the exact mechanism of how this occurs remains to be uncovered.

HisMJP encode for an amino acid ABC transport system. Structural modelling with Phyre2 revealed that HisP shares high structural similarity the crystal structure of an ABC transporter substrate binding protein of *S. pneumoniae* bound to histidine. Cu sequestration within the cytoplasm or periplasm by Cu-binding proteins is one mechanism bacteria employ to subvert Cu toxicity. These protein Cu binding sites are dominated by histidine, cysteine, and methionine residues, with Cu(II) having affinity for histidine (25). Interestingly, GBS are auxotrophic for the amino acid histidine. They overcome this using two different mechanisms; by importing histidine from the environment, or by importing peptides with the aid of specific peptide permeases, which are then subsequently cleaved by intracellular peptidases to their single amino acids (26, 27). This suggests that a GBS Δ*hisMJP* mutant might not have sufficient intracellular histidine to produce proteins rich in histidine residues, either affecting its ability to sequester excess Cu, or to produce yet-to-be characterised proteins essential for growth that require Cu as a cofactor. Evidence of this happening can be seen in our growth experiments and ICP-OES analyses. In our growth experiments, the Δ*hisMJP* mutant reaches a similar CFU and absorbance as compared to the WT after 12 h of growth in THB + 1.5mM Cu. In the nutrient rich THB media, the mutant might be able to overcome its inability to transport histidine into the cell by utilizing extracellular peptides. However, when the Δ*hisMJP* mutant is incubated in minimal CDM + 1.0mM Cu, growth is completely abrogated. Additionally, the Δ*hisMJP* mutant possessed the least amount of total intracellular Cu when subjected Cu stress, which could suggest that the mutant also had less protein-bound Cu within the cell, or that the low levels of Cu were affecting CopA expression/function, which would in turn affect other Cu resistance pathways.

In our study, we also identified several genes (*ackA, ribD, ribE, ribA and ribH*) where an enrichment of insertions resulted in an enhanced Cu tolerance phenotype. To investigate this further, we generated an isogenic mutant in one of these genes (*ribD*), and found that the Δ*ribD* mutant was able to grow better in the presence of 1.5mM Cu as compared to WT in THB media. In GBS, the *ribDEAH* operon encodes a putative riboflavin synthesis pathway. Notably, these genes were down-regulated in response to Zn stress in WT GBS (28), suggesting that riboflavin (or the downstream by-products generated by the pathway) may play a broad role in affecting tolerance to metal ion stress. Interestingly, four other genes (*stp1, yceG. plyB* and *rfaB*) identified in this study also contribute to zinc tolerance (28), suggesting that a common pool of genes might play a role in promoting resistance to metal ion intoxication in GBS.

Establishing the collection of genes that confer GBS resistance to Cu stress dramatically expands our understanding of metal management in this organism, and offers new insight into the diversity of genes that mediate resistance to Cu intoxication in bacteria. For example, genes encoding enzymes for metabolism and cell wall synthesis, regulators, and transporters are critical for GBS to resist Cu stress; future characterization of these will define their effects on the cellular processes that underpin resistance to Cu intoxication.

Bacterial resistance to metal stress is a fitness trait of some pathogens that is used to evade host defence responses (29, 30), and several studies have shown that Cu management contributes to bacterial pathogenicity. For example, *S. pneumoniae* regulates central metabolism in response to metal stress to support its survival (18), and uses CopA for virulence during infection (10). In *E. coli*, Cu-transporting ATPases, including CopA are required for survival in macrophages (31). We recently demonstrated that *copA* contributes to the ability for GBS to colonize and survive in the mammalian host during acute infection (12). Together, these studies and the results of the current work show that Cu management is an important facet of various bacterial pathogens, including GBS in their ability to infect a host. Other observations of bacterial pathogens support a role for Cu management in bacterial virulence in host niches. For example, increased expression of *copY* in *S. pneumoniae* in the lungs of mice was reported (10), and higher Cu levels along with co-incidental up-regulation of *copYAZ* in the blood of mice infected with *S. pyogenes* was reported (11). Characterization of the contributions of the genes of the Cu resistome identified in this current study to GBS virulence will be important to explore potential roles in pathogenesis.

Overall, our application of a highly saturated mutant library combined with deep sequeuncing provides valuable insight into the Cu stress resistome of GBS. Our study identifies a unique collection of genetic targets (including *hisMJP, oafA, yceG, plyB, ribD, rfaB* and *stp1*) that are new to the field of metal detoxification in bacteria and it will be of interest to study their effects towards resistance to metal stress in other pathogens. Together, these findings provide new insight into the repertoire of mechanisms used by GBS to survive Cu stress, and which may be relevant to other bacteria.

## Materials and Methods

### Bacterial strains, plasmids and growth conditions

GBS, *E. coli* and plasmids used are listed in Supplementary Table S2. GBS was routinely grown in Todd-Hewitt Broth (THB) or on TH agar (1.5% w/v). *E. coli* was grown in Lysogeny Broth (LB) or on LB agar. Routine retrospective colony counts were performed by plating dilutions of bacteria on tryptone soya agar containing 5% defibrinated horse blood (Thermo Fisher Scientific). Media were supplemented with antibiotics (spectinomycin (Sp) 100μg/mL; chloramphenicol (Cm) 10 μg/mL), as indicated. Growth assays used 200μL culture volumes in 96-well plates (Greiner) sealed using Breathe-Easy® membranes (Sigma-Aldrich) and measured attenuance (*D*, at 600nm) using a ClarioSTAR multimode plate reader (BMG Labtech) in Well Scan mode using a 3mm 5×5 scan matrix with 5 flashes per scan point and path length correction of 5.88mm, with agitation at 300rpm and recordings taken every 30min. Media for growth assays were THB and a modified Chemically-Defined Medium (CDM) (24) (with 1g/L glucose, 0.11g/L pyruvate and 50mg/L L-cysteine), supplemented with Cu (supplied as CuSO_4_) as indicated. For attenuance baseline correction, control wells without bacteria were included for Cu in media alone.

### DNA extraction and genetic modification of GBS

Plasmid DNA was isolated using miniprep kits (QIAGEN), with modifications for GBS as described elsewhere (32). Mutant strains (Supplementary Table S2) were generated by isogenic gene-deletions, constructed by markerless allelic exchange using pHY304aad9 as described previously (28, 33). Primers are listed in Supplementary Table S3. Mutants were validated by PCR using primers external to the mutation site and DNA sequencing.

### Whole bacterial cell metal content determination

Metal content in cells was determined as described (34). Cultures were prepared essentially as described (12); THB medium was supplemented with 0.5 mM Cu or not supplemented (Ctrl), and following exposure for 2.5h, bacteria were harvested by centrifugation at 4122 x g at 4°C. Cell pellets were washed 3 times in PBS + 5mM EDTA to remove extracellular metals, followed by 3 washes in PBS. Pelleted cells were dried overnight at 80°C and resuspended in 1mL of 32.5% nitric acid and incubated at 95°C for 1h. The metal ion containing supernatant was collected by centrifugation (14,000 x g, 30min) and diluted to a final concentration of 3.25% nitric acid for metal content determination using inductively coupled plasma optical emission spectroscopy (ICP-OES). ICP-OES was carried out on an Agilent 720 ICP-OES with axial torch, OneNeb concentric nebulizer and Agilent single pass glass cyclone spray chamber. The power was 1.4kW with 0.75L/min nebulizer gas, 15L/min plasma gas and 1.5L/min auxiliary gas flow. Cu was analysed at 324.75nm, Zn at 213.85nm, Fe at 259.94nm and Mn at 257.61nm with detection limits at <1.1ppm. The final quantity of each metal was normalised using dry weight biomass of the cell pellet prior to nitric acid digestion, expressed as µg.g^-1^dry weight.

### Transposon Directed Insertion Site Sequencing (TraDIS)

Generation and screening of the 874391:IS*S1* library was performed essentially as previously described (35), with some modifications. Briefly, the pGh9:IS*S1* plasmid (36) (provided by A. Charbonneau *et al*.) was transformed into WT GBS, and successful transformants were selected by growth on THB agar supplemented with 0.5μg/mL Erythromycin (Em). A single colony was picked and grown in 10mL of THB with 0.5μg/mL Em at 28°C overnight. The overnight cultures were incubated at 40°C for 3h to facilitate random transposition of IS*S1* into the bacterial chromosome. Transposon mutants were selected by plating cultures onto THB agar supplemented with Em and growing overnight at 37°C. Pools of the transposon mutants were harvested with a sterile spreader and stored in THB supplemented with 25% glycerol at -80°C. The final library of approximately 480,000 mutants was generated by pooling two independent batches of mutants.

Exposure of the library used approximately 1.9 × 10^8^ bacteria inoculated into 100mL of THB (non-exposed control) or THB supplemented with 1.5mM Cu in THB. The cultures were grown for 12 h at 37°C (shaking), and subsequently, 10mL of culture were removed and washed once with PBS. Genomic DNA was extracted from three cell pellets per condition (prepared as independent biological samples) using the DNeasy UltraClean Microbial Kit (Qiagen) according to the manufacturer’s instructions, except that the cell pellets were incubated with 100 units of mutanolysin and 40mg of RNase A at 37°C for 90min. Genomic DNA was subjected to library preparation as previously described (29), with slight modifications. Briefly, the NEBNext dsDNA fragmentase (New England BioLabs) was used to generate DNA fragments in the range of 200-800bp. An in-house Y-adapter was generated by mixing and incubating adaptor primers 1 and 2 for 2min at 95°C, and chilling the reaction to 20°C by incremental decreases in temperature by 0.1°C. The reaction was placed on ice for 5min, and ice cold ultra-pure water was added to dilute the reaction to 15μM. The Y-adaptor was ligated to the ends of the fragments using the NEBNext Ultra II DNA Library Prep Kit for Illumina (New England BioLabs) according to the manufacturer’s instructions. All adaptor ligated fragments were incubated with *Not*I.HF (New England BioLabs) for 2h at 37°C to deplete plasmid fragments. The digested fragments were PCR amplified as per the protocol outlined in the NEBNext Ultra II DNA Library Prep Kit using a specific IS*S1* primer and reverse indexing primer. DNA quantification was undertaken using a QuBit dsDNA HS Assay Kit (Invitrogen) and purified using AMPure XP magnetic beads (Beckman Coulter). All libraries were pooled and submitted for sequencing on the MiSeq platform at the Australian Centre for Ecogenomics (University of Queensland, Australia). The sequencing data generated from TraDIS libraries were analysed used the Bio-TraDIS scripts (37) on raw demultiplexed sequencing reads. Reads containing the transposon tag (CAGAAAACTTTGCAACAGAACC) were filtered and mapped to the genome of WT GBS 874391 using the bacteria_tradis script with the “--smalt_y 1” and “--smalt_r 0” parameters to ensure accuracy of insertion mapping. Subsequent analysis steps to determine log_2_ fold-change (log_2_FC), false discovery rate (FDR) and P value were carried out with the AlbaTraDIS script (38). To identify genes in 874391 required for resistance to Cu intoxication condition used, we used a stringent criteria of log_2_FC ≤ -2 or ≥ 2, FDR <0.001 and P value <0.05. The TraDIS reads are deposited in the Sequence Read Archive (SRA) under BioProject ID: PRJNA674399.

### Statistical methods

All statistical analyses used GraphPad Prism V8 and are defined in respective Figure Legends. Statistical significance was accepted at P values of ≤0.05.

## Acknowledgments

We gratefully acknowledge Andrew Waller and Amy Charbonneau, Animal Health Trust (Suffolk, UK) for providing pGh9-IS*S1*. We thank Michael Crowley and David Crossman of the Heflin Center for Genomic Science Core Laboratories, University of Alabama at Birmingham (Birmingham, AL) for RNA sequencing. We also thank Lahiru Katupitiya and Dean Gosling for excellent technical assistance. This work was supported by a Project Grant from the National Health and Medical Research Council (NHMRC) Australia (APP1146820 to GCU).

## Supplemental Material

**Supplementary figure S1.**
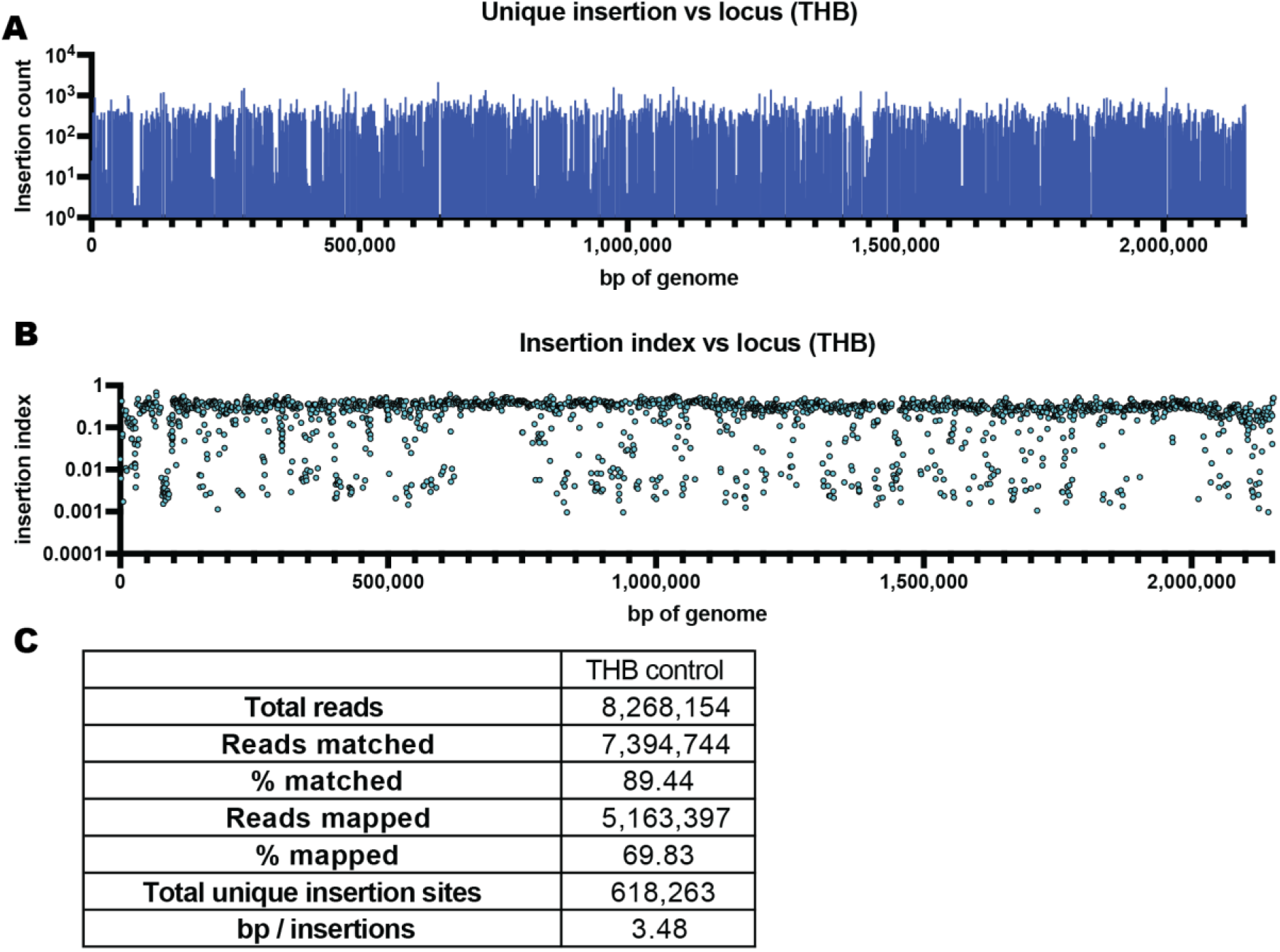
GBS 874391 transposon mutant library. (A) Insertion plot depicting the number of unique transposon insertions within genes along the genome of GBS 874391. Each vertical line represents a single gene, with the height of the line representing the number of unique insertions within the gene. (B) Insertion indices (number of unique insertions / length of gene) of GBS 874391 genes disrupted by IS*S1*. (C) Summary of TraDIS data from sequencing the mutant library recovered after growth in THB. Sequencing reads that matched IS*S1* but did not map to the genome of 874391 mapped to the pGh9:IS*S1* plasmid.

**Supplementary Table 1**. Genes identified in TraDIS screen as significantly under-represented and over-represented.

**Supplementary Table 2**. Bacterial strains and plasmids used in this study.

**Supplementary Table 3**. Primers used in this study.

